# A comparative anatomy of protein crystals: lessons from the automatic processing of 56,000 samples

**DOI:** 10.1101/558023

**Authors:** Olof Svensson, Maciej Gilski, Didier Nurizzo, Matthew W. Bowler

## Abstract

The automatic processing of over 56,000 crystals by the autonomous ESRF beamline MASSIF-1 has provided a data set of crystal characteristics and properties that allows many theoretical proposals and assumptions to be evaluated experimentally.

**Abstract:** The fully automatic processing of crystals of macromolecules has presented a unique opportunity to gather information on the samples that is not usually recorded. This has proved invaluable in improving the sample location, characterisation and data collection algorithms. After operating for four years, MASSIF-1 has now processed over 56,000 samples, gathering information at each stage, from the volume of the crystal to the unit cell dimensions, space group, quality of the data collected and the reasoning behind the decisions made in data collection. This provides an unprecedented opportunity to analyse these data together, providing a detailed landscape of macromolecular crystals and intimate details of their contents and, importantly, how the two are related. The data show that mosaic spread is unrelated to the size or shape of crystals and demonstrate experimentally that diffraction intensities scale in proportion to crystal volume and molecular weight. It is also shown that crystal volume scales inversely with molecular weight. The results set the scene for the development of X-ray crystallography in a changing environment for structural biology.

## 1.0 Introduction

Macromolecular crystallography (MX) has been the primary method for the determination of biological structures over the last 70 years. As such, much effort has been devoted to development of methods to improve the ability to grow crystals, optimise their quality and collect the best possible data from them once they have been placed in an X-ray beam. The end results from these often long and tortuous experiments, structure factors and atomic coordinates, are deposited in one of the earliest examples of a searchable scientific open database, the Protein Data Bank (PDB) (Berman *et al*., 2000). Many studies have used this resource to draw conclusions on the properties of crystals, often with interesting conclusions (Abad-Zapatero, 2012; Berman *et al*., 2013; Berman *et al*., 2015; Robert *et al*., 2017). However, while the database is incredibly useful as a general repository of atomic structures for biologists, it has two fundamental limitations when attempting to draw conclusions on the properties of the crystals themselves. First, the deposited data represent probably the best that were obtained for a sample and were, as such, the result of extensive screening, thereby hiding the potentially thousands of crystals that stand behind the final structure. Second, details of the crystals themselves, such as size, shape, quality variation, data collection strategy etc. are often lost, even if recorded in the primary citation, they are not easy to collate with other data.

Detailed studies have been made on individual systems studying the morphology of the crystals and the packing of the protein, but a general survey across different proteins has never been made, presumably due to the difficulty of gathering such information. More general studies have been made on the PDB itself, producing valuable results in the trends seen in the protein crystals (Berman *et al*., 2013; Berman *et al*., 2015) and also on their physical properties (Robert *et al*., 2017; Bagaria *et al*., 2013) most famously producing the Matthew’s coefficient (Matthews, 1968; Weichenberger & Rupp, 2014). However, both these approaches lack a more general overview of how protein crystal morphology is distributed in general and how this is related to the macromolecule being studied. This is important as it has a direct effect on the requirements of the instrument used to study the crystal (Holton & Frankel, 2010).

The fully autonomous beamline MASSIF-1 at the ESRF (Bowler *et al*., 2015) not only automates the process of sample handling (Nurizzo *et al*., 2016) but also runs complex crystal location, characterisation and decision making routines for every sample processed (Svensson *et al*., 2018; Svensson *et al*., 2015). This level of automation allows a wide range of projects to use the beamline, from those that require extensive screening to find the best diffracting crystal (Li *et al*., 2018; Na *et al*., 2017; Naschberger *et al.;* Sorigué *et al*., 2017; Xu *et al*., 2018) to small molecule fragment screening (Cheeseman *et al*., 2017; Hiruma *et al*., 2017) and experimental phasing at high and low resolutions (Kharde *et al*., 2015; Schulze *et al*., 2018). The routines optimise data collection by centring crystals using X-ray diffraction quality to determine the location of the best volumes to centre (Svensson *et al*., 2015), measuring crystal volumes to dynamically adapt the beam dimeter to match the crystal (Svensson *et al*., 2018) and also to determine the dose the sample can receive before sustaining significant radiation damage (Bowler *et al*., 2016; Svensson *et al*., 2015; Zeldin *et al*., 2013; Bourenkov & Popov, 2010). Samples are then characterised, and optimised data sets collected (Bourenkov & Popov, 2010; Incardona *et al*., 2009), with subsequent autoprocessing of data (Monaco *et al*., 2013; Kabsch, 2010; Vonrhein *et al*., 2011). All of the results from each step of these processes are stored with a unique identification for each sample (Brockhauser *et al*., 2012; Delagenière *et al*., 2011). Crucially, as the samples have been run without any human involvement, the reasons for decisions taken are known and strategies have not been altered before the final data set is collected.

These data have not only allowed individual data collection strategies to be improved but have also improved general strategies by allowing, for example, the most commonly observed crystal dimension to determine the default beam diameter (Svensson *et al*., 2015), improved low resolution data collection strategies or simply to assess the correlation between predicted and obtained resolution (Svensson *et al*., 2018). While these data have proved invaluable in the improvement of the beamline, they also have inherent value, in that they allow the first global survey of crystals of biological macromolecules. Here, we analyse the properties of the 56,459 samples sent to MASSIF-1 between September 2014 and December 2018. The results provide the first general overview of the morphology of crystals of biological macromolecules and how these properties relate to the macromolecule itself and is the first study of its kind in the history of macromolecular crystallography. Together, the results allow many long held assumptions to be tested experimentally and provide a framework to direct the development of future beamline facilities.

## 2.0 Methods

### 2.1 The crystal cohort

MASSIF-1 started taking user samples in September 2014 and has been gathering data on all aspects of these crystals since then. To date, the beamline has processed 56,459 samples from a wide variety of projects and laboratories across Europe and the world (Bowler *et al*., 2016). We believe that this number and distribution of samples represents a reasonable snapshot of crystals for modern structural biology projects. This cohort can therefore form the basis of an analysis that we hope will be generally applicable. The fate of these samples is shown in Table 1.

**Table 1.** The fate of crystals sent to MASSIF-1.

| Stage achieved | Number of samples | Percentage of total |
| --- | --- | --- |
| Samples received at MASSIF-1 | 56,495 | 100 |
| Samples successfully centred | 33,905 | 60 |
| Samples indexed and a strategy calculated | 14,495 | 26 |
| Samples with a processed data set | 16,034 | 28 |

### 2.2 Databases and analysis

All information gathered and used in the automatic location, characterisation and data collection from crystals processed on MASSIF-1 is stored in two databases – one related to the to the sample location, positioning and characterisation processes (Brockhauser *et al*., 2012; Svensson *et al*., 2015) (BES-DB, Support Square, https://supportsquare.io/portfolio/products/) and the second ISPyB (Delagenière *et al*., 2011) records the results of characterisation and data processing. The information for samples contained in each database can be correlated using a data collection ID that is unique to each sample. This allowed us to reconcile the data for samples between databases. A simple python GUI was developed to access data from both databases and store relevant parameters for each sample in *json* format. Analysis of these data was performed using *scipy* and *matplotlib* (Hunter, 2007; Oliphant, 2007).

Crystal dimensions are measured from the X-ray centring routine. The dimensions *x, y* and *z* are the measured crystal width paralell to the spindle axis, the height orthogonal to the spindle axis and the depth orthogonal to the spindle axis 90° away in ω, respectively. The full width half maxima (FWHM) of diffraction signal over images are used to determine crystal dimensions. Over sampling means that the minimum distance that can be measured using the 50 μm diameter beam is 25 μm, dimensions smaller than this are determined using the smaller beam apertures. As the automesh algorithm (Svensson *et al*., 2015) will place the sample mount at either the smallest or widest orientation of the mount in ω, depending on wheather single or multiple data collections are requested (Svensson *et al*., 2018), we are confident that in most cases the dimensions measured will be consistent with the orientation of the crystals as they tend to lie paralell to the mount. This would reduce the overestimation of sample height and depth, for instance if a plate shaped crystal was presented at an angle. All samples are assumed to be cuboid.

Unit cell volumes were calculated from the dimensions obtained during indexing using the following equation:

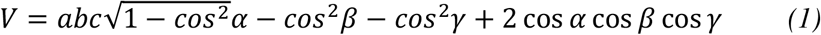

The molecular weight (kDa) of the entity in the asymmetric unit was estimated using the following equation:

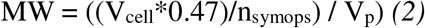

where a solvent content of 47% was assumed (PDB average), n_symops_ is the number of symmetry operators determined from the point group and V_p_ is derived from the partial specific volume for globular proteins of 0.73 cm^3^ g^−1^ (Harpaz *et al*., 1994), here expressed as 1210 Å^3^ kDa^−1^. These assumptions are reasonable, but will lead to some overestimates of molecular weight for smaller proteins and an under estimation for larger proteins, where the solvent contents may be significantly different to 47%. The molecular weight will also be inaccurate if the crystal is incorrectly indexed, particularly if the triclinic point group is incorrectly selected. The process was verified by comparing 7 proteins with known molecular weights to the values calculated using these methods (see Table S1) and the average is the same as that of the PDB (Berman *et al*., 2013). A histogram of the molecular weight distribution is shown in Figure S1.

## 3.0 Results and Discussion

### 3.1 The size and shape of protein crystals

Accurately determining the dimensions, and therefore volumes, of a protein crystal is primarily important in determining the dose that the crystal can absorb before significant radiation damage (Bourenkov & Popov, 2010; Bowler *et al*., 2016; Svensson *et al*., 2015; Zeldin *et al*., 2013) and in determining the diameter of the beam that should be used to maximise the signal to noise ratio (Evans *et al*., 2011; Svensson *et al*., 2018). However, the gathering of volumetric data has many other potential uses, not least being able to correlate crystal size to quality for individual projects. The data collected for all samples processed on MASSIF-1 provide an opportunity, for the first time, to define the broad distribution of protein crystal dimensions. Here, we show volumes in cubic millimetres. This can be difficult to convert into real world quantities so a comparison between units is shown in Table 2.

**Table 2.**
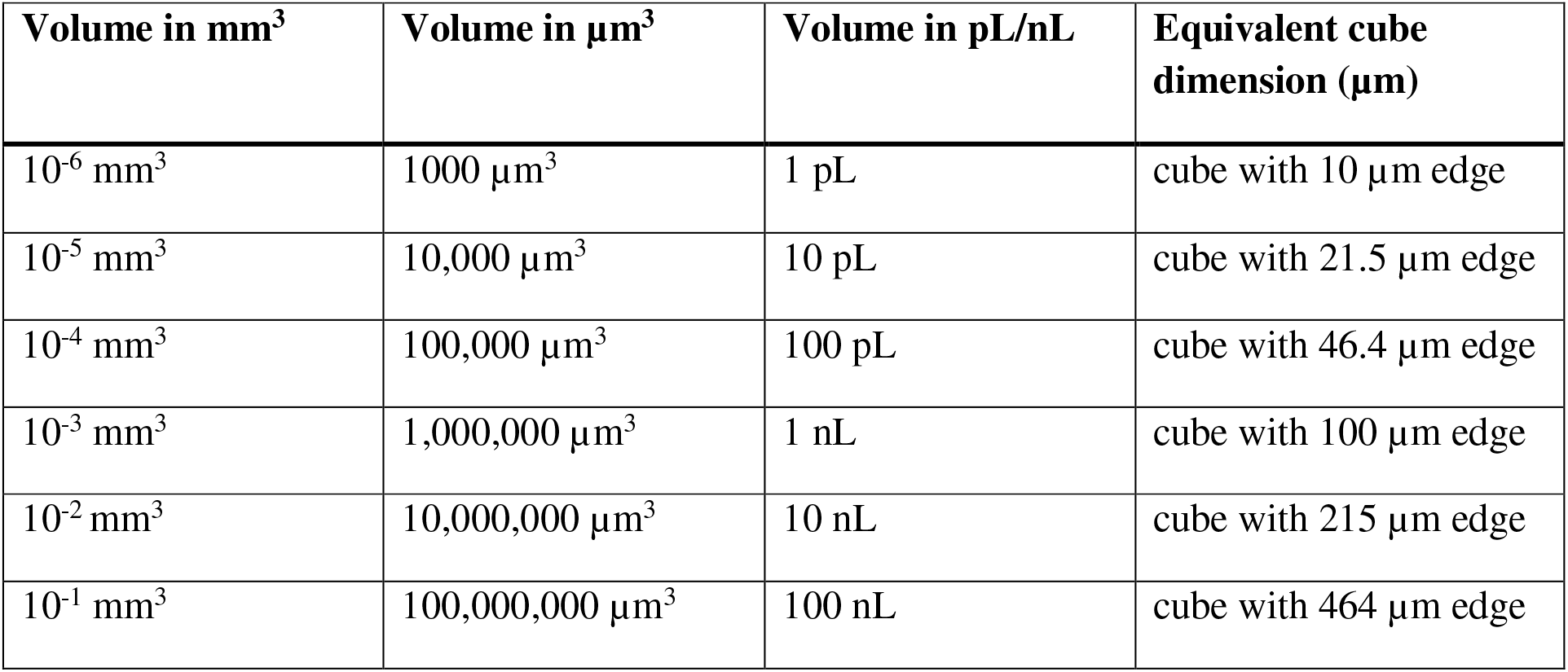
Volume guide

To our knowledge, there has never been a general study of protein crystal volumes. While there are some careful studies of individual proteins (Frey *et al*., 1991; Joachim & Markus, 2015; Liu *et al*., 2013; Mayans & Wilmanns, 1999) we have no data for the general distribution. Programs that account for crystal volume when computing absorbed doses, such as *BEST* (Bourenkov & Popov, 2010; Popov & Bourenkov, 2003) and *RADDOSE3D* (Zeldin *et al*., 2013) are extremely useful when users input the correct crystal dimensions. However, a default volume, a cube with sides of 100 μm, is used in the absence of measured dimensions. This value originated in the early days of MX (Helliwell, 1984), but does it relate but does it relate to the reality of protein crystals today?

The distribution of measured crystal volumes is shown in Figure 1. The mean volume of 0.002303221 mm^3^ (2,303,221 μm^3^), is the same volume as a cube with edges of 132 μm. While the majority of crystals are smaller than this average, the distribution is lognormal and the mode volume is 0.000020209 mm^3^ (20,209 μm^3^, a cube with edges of 27 μm), it does seem to validate the choice of the default average crystal volume used. However, volumetric data alone hide an important factor – morphology. The best way to demonstrate the relationship between a shape and its volume is the surface area to volume ratio. Here, we have plotted the surface area against the volume (Figure 2). The plot shows most crystals have a surface area greater than that expected for a cube with the crystals with very large volumes being more cuboid. What is important is that many crystals that have a large volume (e.g 0.01 mm^3^) have shapes that are best matched by a cylinder of 40 μm diameter or a plate 50 μm thin (Figure 2, magnified panel). This reflects the distribution of measured dimensions, which have a modal value of around 50 μm (Bowler *et al*., 2016; Svensson *et al*., 2015). This implies that having X-ray beams larger than 100 μm will have limited returns and most crystals will require diameters of 10 to 50 μm with, of course, the possibility to collect from multiple volumes in plate or needle crystals.

**Figure 1.**
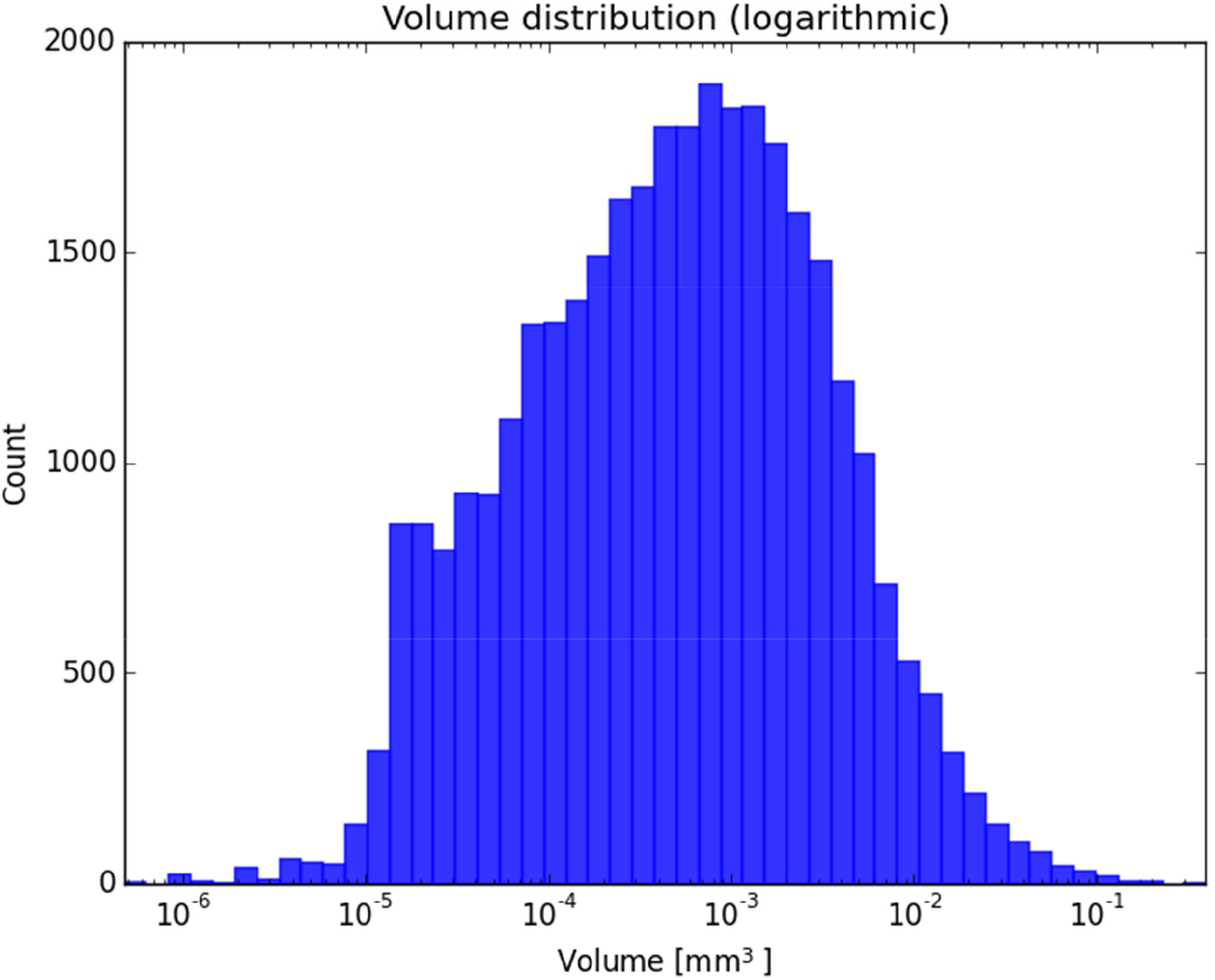
Histogram of crystal volumes. The histogram shows the distribution of crystal volumes measured on MASSIF-1. Note logarithmic scale, N=33,905.

**Figure 2.**
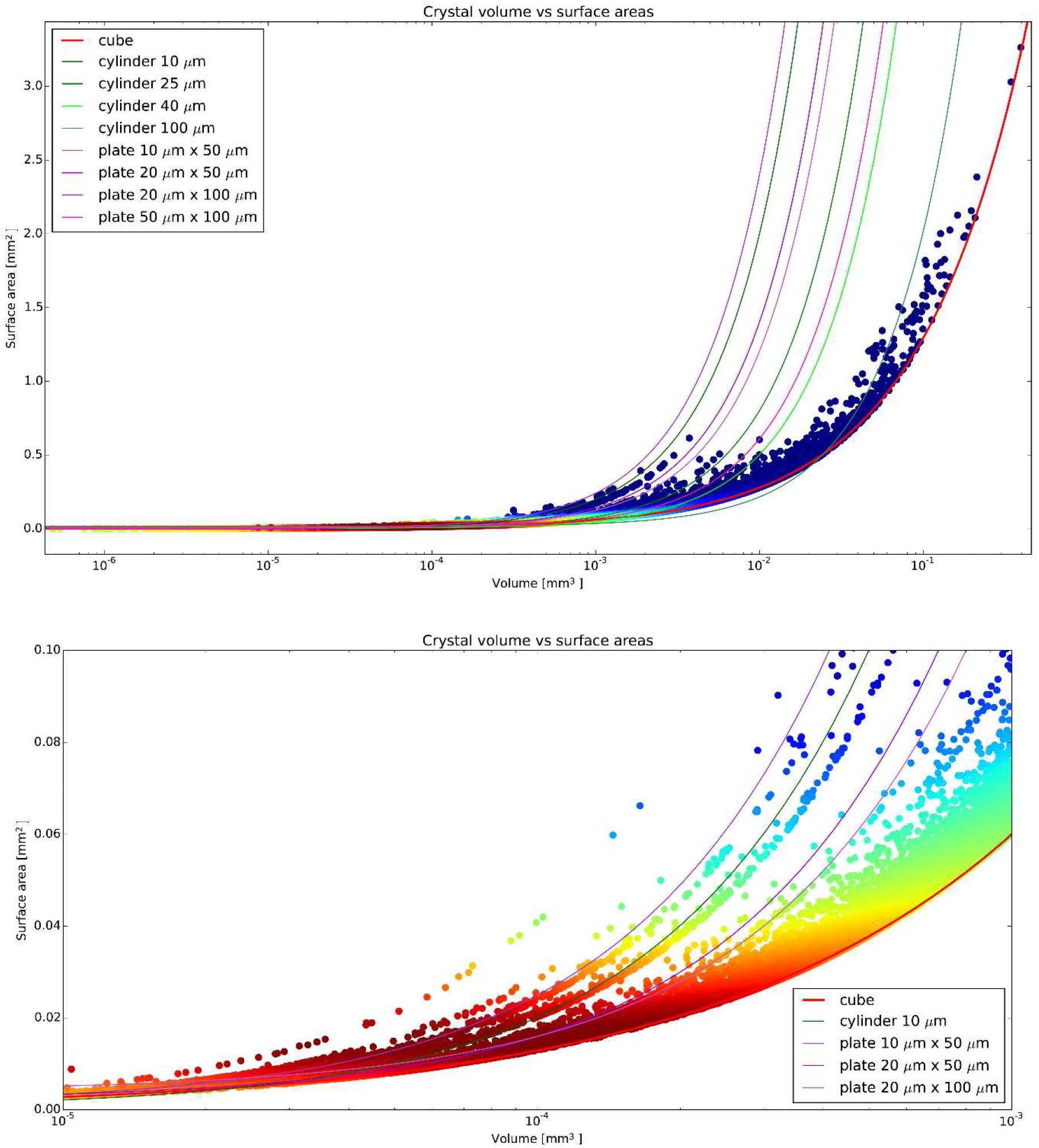
Crystal surface area to volume ratio. The relationship between surface area and volume is shown with lines representing the curves described by different shapes and points coloured by the kernel density estimate (KDE). N=33,905. The lower panel shows a magnified area with the highest counts. Most crystals are small and have one dimension that is in the 10 to 30 μm range.

### 3.2 The internal properties

The recording of volumetric data along with the results from data collection now allows us to test certain maxims within the MX community. It is generally accepted that larger crystals will be more difficult to cryo-cool, for example. The mosaic spread of a crystal has been demonstrated to be closely correlated to effectiveness of the cooling protocol employed (Mitchell & Garman, 1994; Kriminski *et al*., 2002) and we have used this measure to relate to volumetric data. The mosaic spread value used here is the *MOSFLM* estimated value from four characterisation images (Leslie, 2006) and is used as it is the only value calculated in the same manner for all samples. This value can be higher than that calculated by, for example XDS (Powell *et al*., 2017; Kabsch, 2010), but will be consistent and allow trends to be discerned. Plotting crystal volume against mosaic spread (Figure 3A) does not show a correlation, in fact there is a weak negative correlation (Spearman R = −0.26). As the cooling rate is an important factor in cryo-cooling (Garman, 1999; Teng & Moffat, 1998) perhaps the surface area to volume ratio (S/V) is more important? Crystals with a higher S/V should cool faster. Plotting S/V against mosaic spread (Figure 3B) again does not show that crystals that could potentially cool more rapidly have lower mosaic spread values and again there is a weak negative correlation (Spearman R = 0.23). If the mosaic spread is independent of crystal shape and size, is it more closely related to the entity crystallised? Plotting the molecular weight against mosaic spread (Figure 3C) does seem to point to a trend to higher mosaic spread values for larger macromolecules, but again the correlation is weak (Spearman R = 0.2). Is then the order of the crystal more dependent on the entity crystallised? While the correlation shown here is weak, small molecule crystallographers have observed that smaller crystals have higher mosaic spread values (Andrews *et al*., 1987; Andrews *et al*., 1988; Papiz *et al*., 1990) proposing that crystal growth could be limited by the disorder in the crystal. While the variation in mosaic spread values we have measured here is most likely attributable to preparation of the crystals, it should not be ruled out that mosaic spread could be an inherent property of the crystal present in the drop. This seems to be supported by the lower mosaic spread values for more cuboid crystals (Figure 3B) implying that well ordered growth in all lattice directions is a better predictor of lower mosaic spread. Several studies have shown that mosaic spread can be high at room temperature and reduced via controlled dehydration (Bowler *et al*., 2006; Sanchez-Weatherby *et al*., 2009; Russi *et al*., 2011; Amunts *et al*., 2007; Kiefersauer *et al*., 2000) indicating that mosaic spread can already be high before cryo-cooling. This seems to counter received wisdom that protein crystals tend to have lower mosaic spread values at room temperature (Garman, 1999) but only a few systematic studies have been made (Fischer *et al*., 2015; Low *et al*., 1966; Juers & Matthews, 2001) making this a difficult conclusion to support as generally applicable.

**Figure 3.**
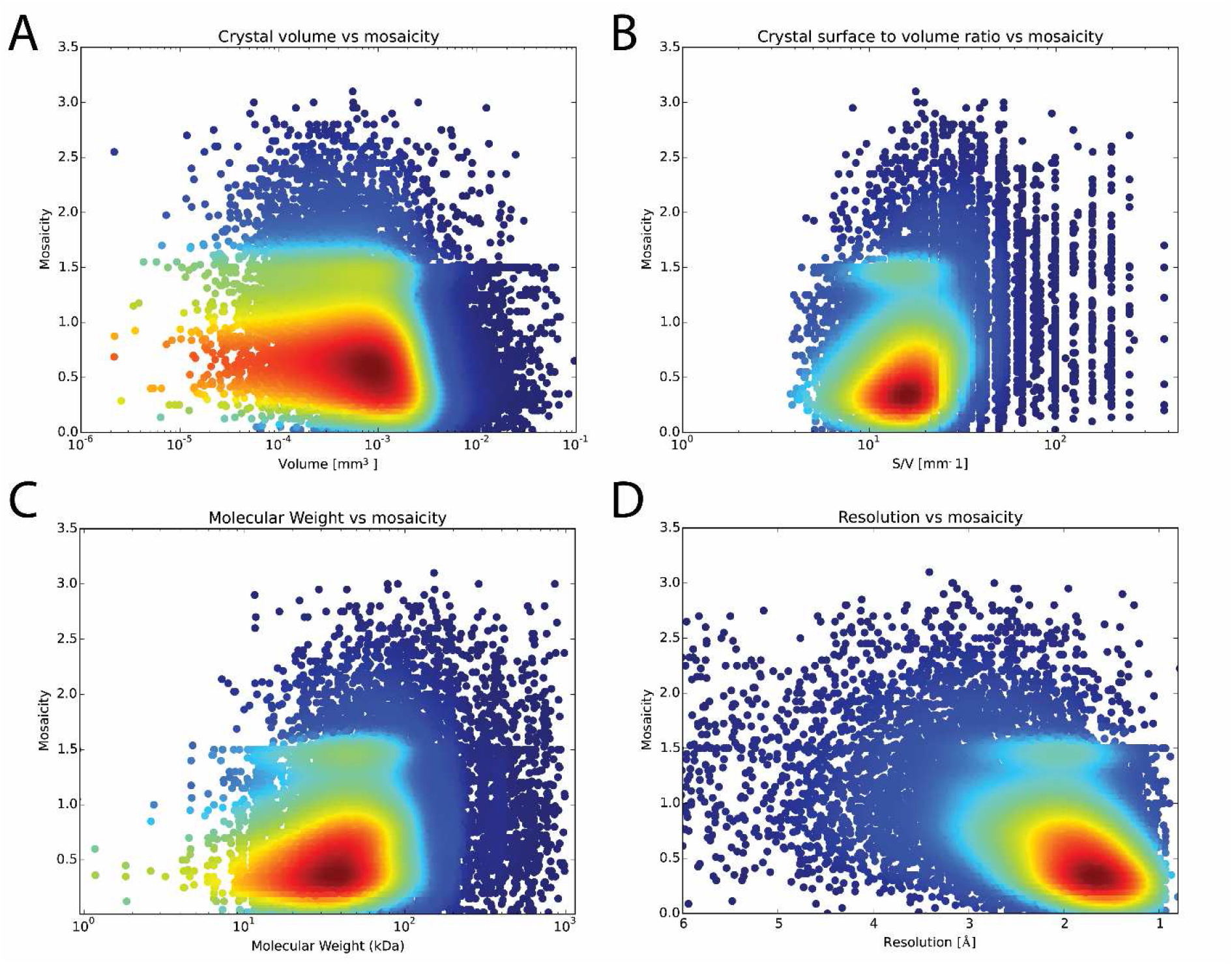
What parameters correlate with mosaic spread? Mosaic spread values are plotted against crystal volume, Spearman R = −0.26 (**A**), surface area to volume ratio, Spearman R = 0.23 (**B**), molecular weight of the entity crystallised, Spearman R = 0.2 (**C**) and the resolution cut-off of the processed data set, Spearman R = 0.44 (**D**). Data points are coloured by KDE, N=14,234 for each panel.

What practical implications does this have? It seems to be clear is that crystals should not be selected based on their size and shape. It is therefore probably more important to focus on careful crystal handling to minimise mosaic spread through choice of cryo-protectant, soaking protocol and speed of cooling (Garman, 1999; Warkentin *et al*., 2006). It is worth spending time obtaining lower mosaic spreads, when plotted against resolution there is a good correlation (Spearman R = 0.44, Figure 3D).

Another important parameter is the variation of diffraction quality within a crystal. It has been shown that the different crystal volumes can vary widely leading to significantly better data sets from the more ordered regions (Bowler *et al*., 2010; Thompson *et al*., 2018) but how common is it for crystals to diffract heterogeneously? Previous work has defined a measure of diffraction variability within crystals that was demonstrated on 19 test samples (Bowler & Bowler, 2014). The measures, *V_1_* and *V_2_*, define variability as the variance in diffraction quality over the mean squared, and the peak value over the mean, respectively. A simple model defines the ratio *N* giving an idea of the proportion of the crystal that varies and how great the difference in diffraction quality is. At the time of the initial study, the total integrated signal of the images collected during a mesh scan was used to define diffraction quality. Since MASSIF-1 started, the measure is the *Dozor* score (Svensson *et al*., 2015; Melnikov *et al*., 2018). *Dozor* determines the distribution of background intensity, azimuthally averages the spot intensities and removes areas showing ice or salt diffraction. The mean intensity of Bragg spots against resolution over background is then determined and used to create a score of quality. A plot of *V_1_* against *V_2_* with various ratios N, from all mesh scans performed on MASSIF-1 using the *dozor* score as metric, is shown in Figure 4A. From the plot it can be seen that most crystals are quite homogenous, displaying ratios *N* below 5. However, there are a large number of observations where the diffraction quality varies enormously, with peaks 4 to 6 times above the average (it should be noted that the *dozor* score only varies between 10-20% within images of a dataset from a single position). Several lines can be seen defined by data points: these describe lines of *N*=0.5, 1 and 2 and arise from small crystals that have been probed in only 2 or 3 positions. Two positions can only give *N* =1.0 and either *N* =0.5 or 2.0 for 3 positions. The vast majority of these results come from scanning with a 50 μm beam diameter, smaller or larger diameters are only used when specifically requested by users (Svensson *et al*., 2018; Svensson *et al*., 2015). How does the variability relate to other characteristics? When compared to mosaic spread there is no correlation (Figure 4B) nor with the molecular weight of the entity crystallised (Figure 4C). When compared to the resolution of the final data set there is a weak correlation for higher resolution for a higher ratio *N* (Figure 4D). As the final data set is collected from a single position, with the beam diameter adapted to the best region, it is perhaps not entirely surprising that good data can still be collected from a good region of a variable crystal than a selected region of a homogenous crystal and it is clear that heterogeneous quality does not prevent the collection of a good quality data set if the correct strategy is employed. Given the significant variation in quality observed, the ultimate strategy for scanning would be to use the smallest possible beam to probe variation and then adapt the diameter to match the size of the best volume determined.

**Figure 4.**
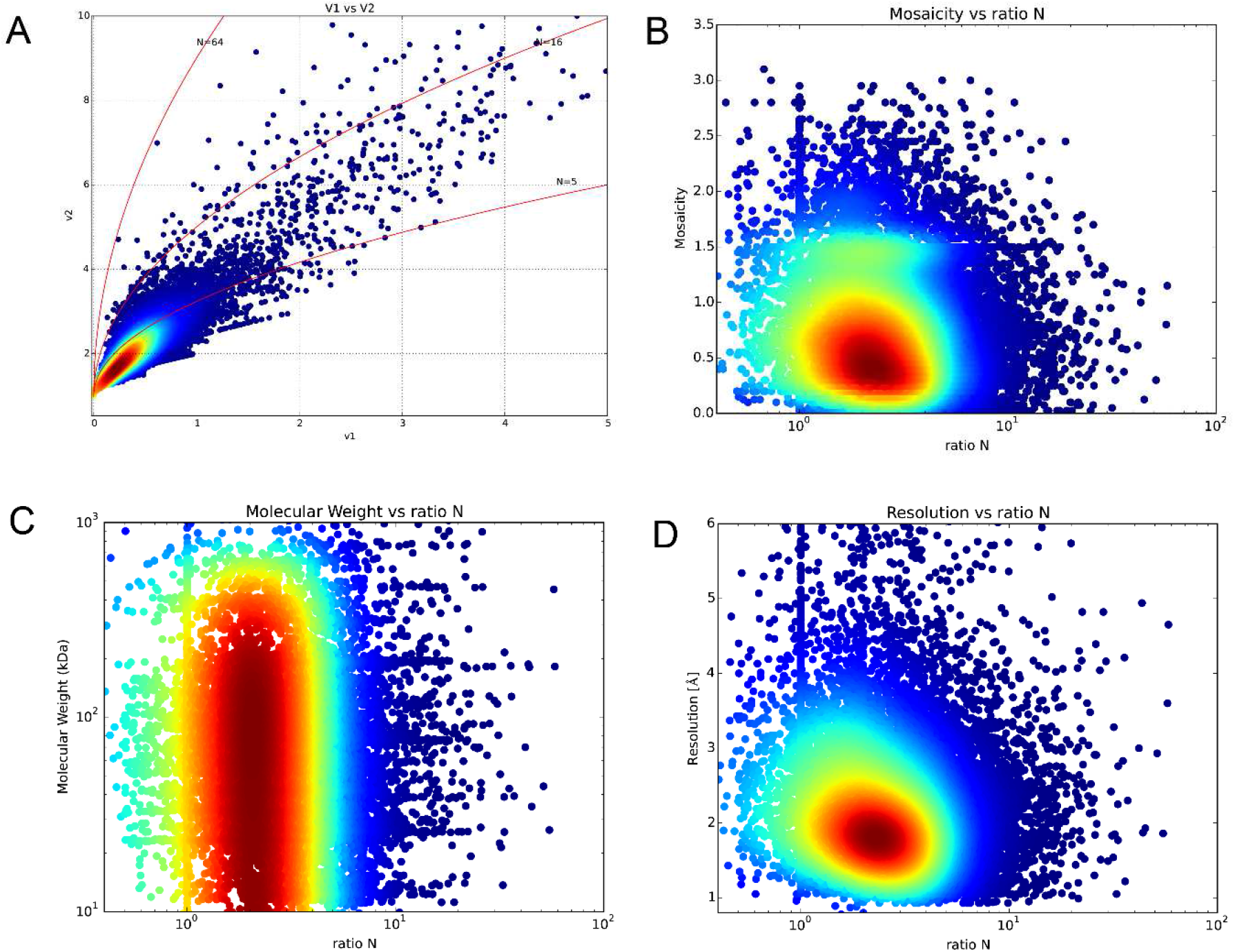
The variation of quality within crystals. **A**. Comparison of variability measures of crystals. Values of *V_1_* and *V_2_* are plotted against each other and coloured by KDE. Lines show the values obtained for various ratios *N* between positions at increasing differences in diffraction power (see (Bowler & Bowler, 2014) for an explanation of the model). The red line representing the ratio *N* = 5 is a reasonable cut-off between variable and homogenous diffraction within crystals. N=15,864. **B**. Mosaic spread values plotted against ratios *N*. Greater variability within a crystal is not related to higher mosaic spread values, Spearman R = −0.13, N=13,780. **C**. The molecular weight has no effect on the degree of variability, Spearman R = −0.06, N=15,188. **D**. Obtained resolution against ratios *N*, higher variability is weakly correlated to higher resolution, Spearman R = −1.9, N=15,188

### 3.3 Relationship between crystal volume, molecular weight of the protein and resolution

The scattering power of a crystal depends on the number of unit cells that can be illuminated in the beam, meaning that the volume of both the crystal and the unit cell, as well as the properties of the molecule, are critical to a successful experiment (Holton, 2009). Practically, this means that the larger the molecule being studied, the larger its B factor, and the higher the required resolution, the larger the crystal will have to be. How does this relate to the actual measured values of crystals that yielded data sets on MASSIF-1? First, the number of unit cells can be plotted against the crystal volume (Figure 5). While not surprising, it is informative to see how the number ranges across projects that make it through to indexing – 10^12^ unit cells is most common (mode) and the smallest number that led to a processed data set was 2.3 x 10^8^ unit cells from a crystal 10 x 12 x 20 μm^3^ in size, 15 μm beam diameter, *P*6 *a* = *b*= 85.94 Å, c = 201.45 Å, α = β = 90°, γ = 120°. A thorough theoretical treatment for this relationship has been demonstrated (Holton & Frankel, 2010; Holton, 2009) defining the minimum crystal volume required under ideal conditions. The theoretical relationship has been extremely useful for predicting the requirements and limits of new facilities and for providing a target for beamlines to aim for when optimising experiments (Grimes *et al*., 2018). Does the relationship hold experimentally over a large number of different samples? Figure 6 shows the resolution obtained plotted against crystal volume coloured by the molecular weight of the molecule crystallised. The theoretical relationship for the minimum required crystal volume is also plotted. The lines describe *equation 16* from (Holton & Frankel, 2010) assuming a Nave-Hill effect (photoelectron escape) of 1, as all crystals are larger than 1 μm^3^, and 100 photons per *hkl* that approximates the experimental conditions on MASSIF-1. The agreement between the theoretical curves and observed samples is remarkable (Figure 6) with most crystals remaining above the minimum volume predicted for a given resolution and molecular weight. The curves represent specific molecular weights and B factors, which can vary enormously, and should be taken as a guide to where these values lie. The observation confirms the theoretical treatment as excellent and it clearly defines the standards that beamlines should be aiming for.

**Figure 5.**
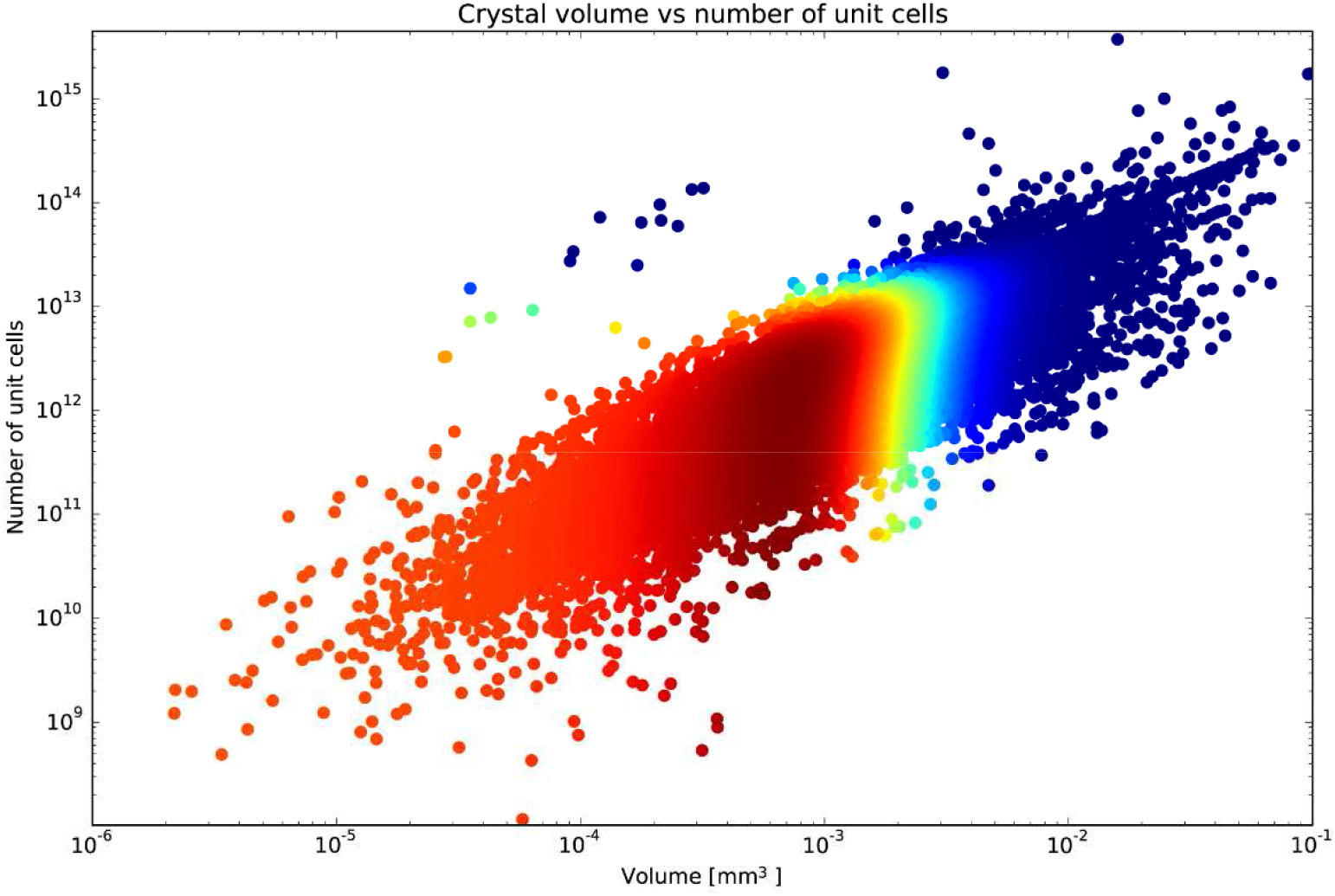
Number of unit cells plotted against crystal volume. There is a large spread across 3 orders of magnitude in the number of unit cells in the highest count bins. The smallest number of unit cells in a crystal the yielded a data set was 2.3 x 10^8^. Data points are coloured by KDE, N=15,905.

**Figure 6.**
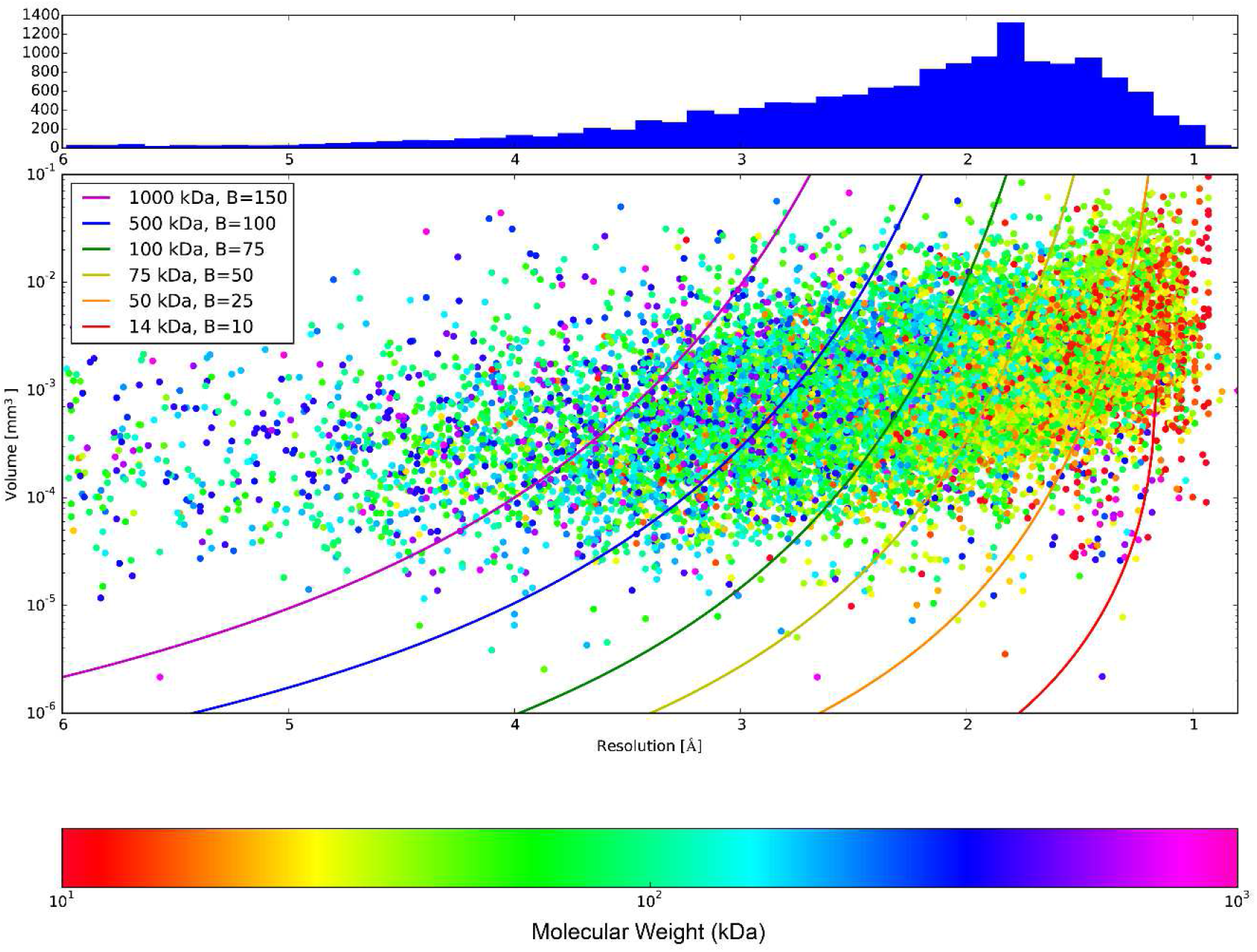
The relationship between crystal volume, obtained resolution and molecular weight. The crystal volume is plotted against the highest resolution cut-off from autoprocessing and coloured by the molecular weight of the entity crystallised, note logarithmic scales for volume and molecular weight. Lines describe the minimum crystal volume required for a certain resolution given the parameters shown assuming a Nave-Hill effect (photoelectron escape) of 1, as all crystals are larger than 1 μm^3^, and 100 photons per *hkl* (equation 16 from (Holton & Frankel, 2010)). The lines indicate a specific molecular weight and B factor and should be taken as a guide to the rough range described. The wavelength on MASSIF-1 is fixed and the beam diameter is altered to match the crystal. The histogram shows the distribution of resolutions obtained in the scatter plot. N=15,241.

A more informative plot shows the molecular weight of the molecule crystallised plotted against the volume of the crystal coloured by the final resolution obtained from the data set (Figure 7). The most striking observation is that, on average, the larger the molecule the smaller the crystal. This is rather unfortunate given the dependence on volume for a given resolution. It is also interesting as a distribution of the molecules studied – there is a clear drop off after ~200 kDa, showing the current range of samples studied by MX. This cut-off is significant when considering the role of synchrotron beamlines in the future of structural biology. With technological advances in cryo-electron microscopy (cryo-EM) allowing structure determination at medium resolutions for very large complexes (Subramaniam *et al*., 2016), X-ray crystallography will soon no longer be the method of choice for systems over ~120 kDa. While analysing proteins below this molecular weight is possible by cryo-EM (Khoshouei *et al*., 2017) it remains extremely difficult, requiring an excellent sample and not yet obtaining the same resolutions and speeds of data acquisition as a modern synchrotron beamline. Figure 7 demonstrates that the two techniques remain highly complementary, as molecular weight increases, crystal volume and resolution tend to decrease making structure determination by X-ray crystallography harder. This trend is inverted for Cryo-EM, meaning that X-ray crystallography can concentrate on the <100 kDa macromolecules providing high data throughput and resolution with cryo-EM working on the >100 kDa region where resolution will be equivalent, or higher, and experiments less difficult. Most crystals in this molecular weight range lie in the 10^−4^ to 10^−2^ mm^3^ range and would require X-ray beams from 30 to 100 μm in diameter.

**Figure 7.**
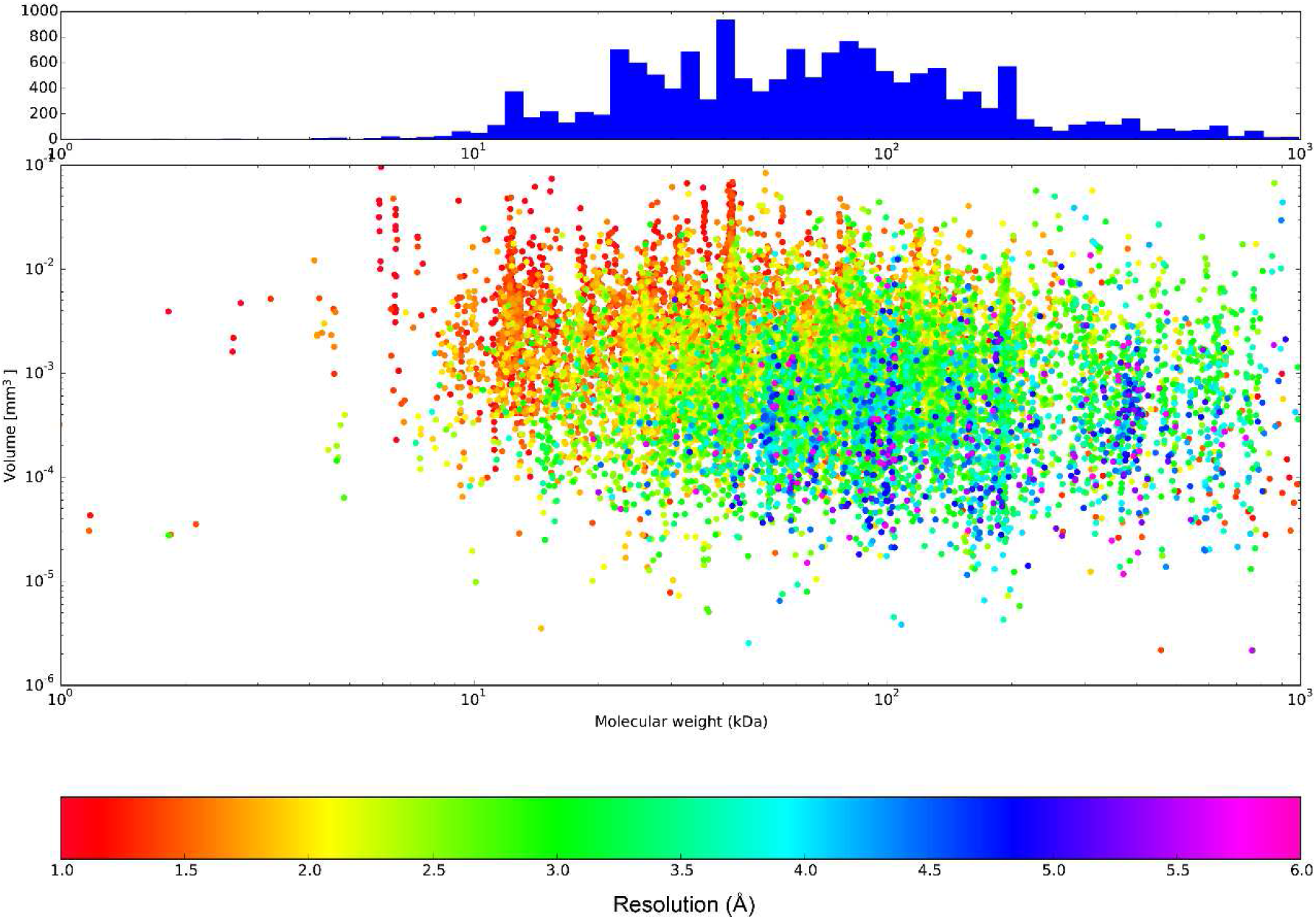
The relationship between crystal volume, molecular weight and obtained resolution. The crystal volume is plotted against the molecular weight of the entity crystallised coloured by the highest resolution cut-off from autoprocessing, note logarithmic scales for volume and molecular weight. The histogram shows the distribution of molecular weight in the scatter plot. N=15,241.

## 4.0 Conclusion

The fully automatic collection of data has allowed the study and comparison of the physical and molecular properties of a wide range of crystals and their constituent macromolecules for the first time. The results provide an overview of how size and shape is distributed over these crystals to a high level of precision, superseding assumptions based on individual systems. Coupling these data to the internal properties of these crystals from the processing of derived data sets we can start to challenge assumptions and theoretical treatments. Have our assumptions been correct? Regarding crystal shapes and sizes, many have been wrong. These data demonstrate that an ‘average’ crystal is more likely to be a plate or needle with a minimum dimension in the 30 to 50 μm range. The expectation that smaller or flatter crystals will cool better also seems to be incorrect and, as is so common in experimentation, sample handling and preparation is probably much more important. It is also reassuring that the theory behind diffraction and radiation damage holds extremely well in the real world.

How can these data be used? Globally, they can guide the development of future facilities demonstrating that a range of beam diameters are required, even if larger complexes will be increasingly top sliced by cryo-EM, and that the theoretical limits should be strived for in an experiment station. It is clear that microfocus beamlines will still be needed and that also beamlines with larger focal spots should not be neglected. Further insights could also be gained by linking the data collected here to crystallisation databases from highly automated facilities (Ng *et al*., 2016; Shaw Stewart & Mueller-Dieckmann, 2014) providing a further link between crystallogenesis and the final result. This link could inform the crystallisation laboratory on the highest quality data as well as volume data and how these relate to the conditions that produced them. Additionally, further information on crystallisation techniques, handling cryoprotection protocols linked to the information presented here would be very valuable (Newman *et al*., 2012). Studies such as this have been limited to the PDB, and while highly informative (Abrahams & Newman, 2019), no data are available on the crystals or screening required to get the result. More specifically, the data gathered can help individual projects by informing on the spread of volumes and how these relate to data quality potentially improving data collection strategies. The analysis of these data has already improved the operation of MASSIF-1 (Svensson *et al*., 2018; Svensson *et al*., 2015) but could this go further? Recent developments in machine learning could be applied to all the data collected and may help to improve data collection strategies. Looking more closely with more data than has previously been available, questions such as: when is a helical or multi-position data collection better than single position? Can specific strategies, such as SAD, be improved? could be answered. The analysis presented here has only started to delve into the data and we hope that modern data science techniques could help further improve the measurement of diffraction data from protein crystals.

## Acknowledgements

We would like to thank all the users of MASSIF-1 over the years for trusting the beamline to physically handle and make data collection decisions for their precious samples. These samples have made the beamline the success it is and have provided the data on which this study is based.

---

**Table S1.** Verification of calculated molecular weights compared to known samples.

| Acronym | Actual MW (kDa) | Calculated MW (kDa) |
| --- | --- | --- |
| Lyso | 14.2 | 12 |
| RICK | 34.85 | 48 |
| CRU | 41.31 | 43 |
| Erk2 | 42.24 | 41 |
| DPP | 162 | 200 |
| TbrATP | 922.6 | 800 |
| GroEL | 910 | 700 |

**Figure S1.**
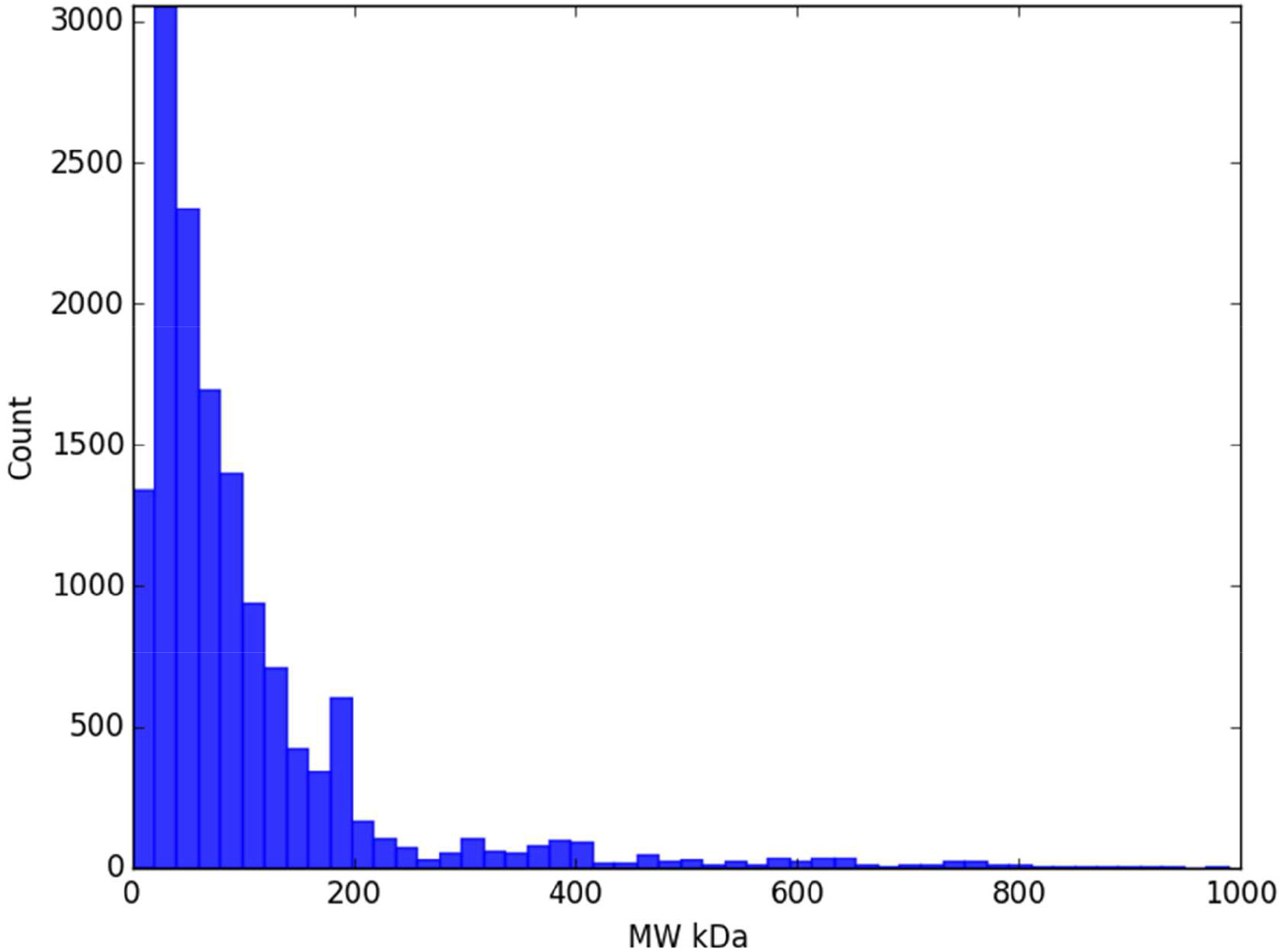
Histogram of molecular weights calculated from samples processed on MASSIF-1.

